# Endogenous Nuclear Desmin Associates with the Nuclear Envelope in Human Skeletal Muscle Cells

**DOI:** 10.64898/2026.09.17.752464

**Authors:** Nilufer Duz, Ahmet Avcı, Sevval Sahin, Mija Kastelic, Sureyya Ozcan, Pervin Dincer

## Abstract

Desmin is the major intermediate filament protein of muscle, yet its potential nuclear functions remain poorly understood. Here, we investigated the nuclear localization and organization of endogenous desmin in human skeletal muscle cells using quantitative confocal microscopy, biochemical fractionation, targeted MRM/SRM proteomics, and computational structural modeling. Desmin localized to the nuclear compartment and exhibited significantly stronger colocalization with emerin and lamin A/C than with lamin B1. Nuclear fractionation and targeted proteomics independently confirmed endogenous nuclear desmin, while protein–protein docking predicted structurally plausible interactions with all three nuclear envelope proteins, supporting a context-dependent nuclear interaction network. Together, these findings identify the nuclear envelope as a functional platform for desmin and suggest that nuclear desmin contributes to nucleo-cytoskeletal communication and the organization of mechanically responsive nuclear architecture during myogenesis. This work expands the current understanding of desmin biology and provides a framework for investigating its role in desmin-related muscle disease.

## INTRODUCTION

The nucleus is a highly organized organelle whose structure and function depend on continuous communication with the cytoplasm. This communication is coordinated by the nuclear envelope, nuclear pore complexes, and the nuclear lamina, which together regulate nucleocytoplasmic transport, nuclear mechanics, chromatin organization, and gene expression. Increasing evidence indicates that, in addition to soluble signaling molecules and transcription factors, several cytoskeletal proteins transiently localize to the nucleus, where they participate in diverse processes including transcriptional regulation, DNA repair, and mechanotransduction. These observations suggest that the nucleus is dynamically influenced by proteins traditionally regarded as exclusively cytoplasmic.^1–8^

Among cytoskeletal proteins, intermediate filaments have emerged as important regulators of nuclear organization. Besides the nuclear lamins, several cytoplasmic intermediate filament proteins, including vimentin and keratins, have been detected within the nucleus under specific physiological or pathological conditions, where they have been implicated in chromatin organization, stress responses, and regulation of gene expression. Nevertheless, the mechanisms governing their nuclear localization and their molecular functions remain poorly understood. These findings raise the possibility that additional intermediate filament proteins may possess previously unrecognized nuclear functions.

Desmin is the principal type III intermediate filament protein of skeletal, cardiac, and smooth muscle cells, where it maintains cellular architecture by mechanically linking myofibrils, mitochondria, the sarcolemma, and the nucleus. Although traditionally regarded as a cytoplasmic structural protein, several independent observations challenge this view. Nuclear localization of desmin has been reported in cultured cells, during cardiomyocyte differentiation, and following interactions with nuclear components, suggesting that a subset of desmin enters the nuclear compartment. The biological importance of these observations is underscored by a homozygous *DES* splice-site mutation (c.1289-2A*>*G) within the lamin-binding region that causes limb-girdle muscular dystrophy R2/myofibrillar myopathy. Despite preserved desmin filament organization, this mutation disrupts desmin–Lamin B interactions, while patient-derived cells exhibit impaired myogenic differentiation, indicating that defective nuclear functions of desmin may contribute to disease pathogenesis. Recently, we demonstrated that desmin contains two functional nuclear localization signals and undergoes active import through the *α/β*-dependent pathway, establishing that its nuclear localization is a regulated process rather than passive diffusion. However, despite growing evidence for nuclear desmin, the precise subnuclear localization of endogenous desmin and its molecular interactions with nuclear envelope proteins remain unknown. ^22^

To address these unresolved questions, we investigated whether endogenous desmin localizes to the nuclear compartment and whether it preferentially associates with specific components of the nuclear envelope in human skeletal myoblasts. We combined high-resolution confocal microscopy with quantitative colocalization analyses, biochemical subcellular fractionation, targeted MRM/SRM proteomics, and protein–protein docking to define the subnuclear distribution of desmin and examine its relationship with Lamin B1, lamin A/C, and emerin. Unlike previous studies that primarily established the existence or transport mechanism of nuclear desmin, our approach enabled quantitative characterization of its endogenous nuclear localization and provided complementary biochemical and structural evidence supporting its association with distinct nuclear envelope proteins. Together, these findings establish a molecular framework for understanding the nuclear organization of desmin and provide new insight into how disruption of these interactions may contribute to skeletal muscle differentiation and desmin-associated myopathies.

## RESULTS

### Desmin preferentially associates with lamin A/C and emerin at the nuclear envelope

To investigate the spatial relationship between desmin and nuclear envelope proteins, we performed confocal immunofluorescence imaging followed by quantitative fluorescence signal overlap analyses. Desmin exhibited a prominent perinuclear distribution pattern and partial spatial overlap with Lamin A/C, Lamin B, and emerin signals (Figure 1). Signal overlap maps generated from deconvolved z-stack images revealed enrichment of desmin-associated fluorescence at the nuclear periphery. Notably, nuclear desmin localization was heterogeneous, as colocalization with nuclear envelope proteins was detected only in a subset of nuclei within the same imaging field. A similar heterogeneous pattern was observed in multinucleated myotubes, where individual nuclei within the same myotube displayed variable degrees of desmin colocalization. Representative colocalization maps from consecutive optical sections of the same imaging field, together with the corresponding Pearson’s correlation coefficients, provide a qualitative and quantitative visualization of the z-dependent changes in desmin–emerin colocalization that underlie the analyses shown in Figure 2A (Supplementary Figure S1).

**Figure 1.**
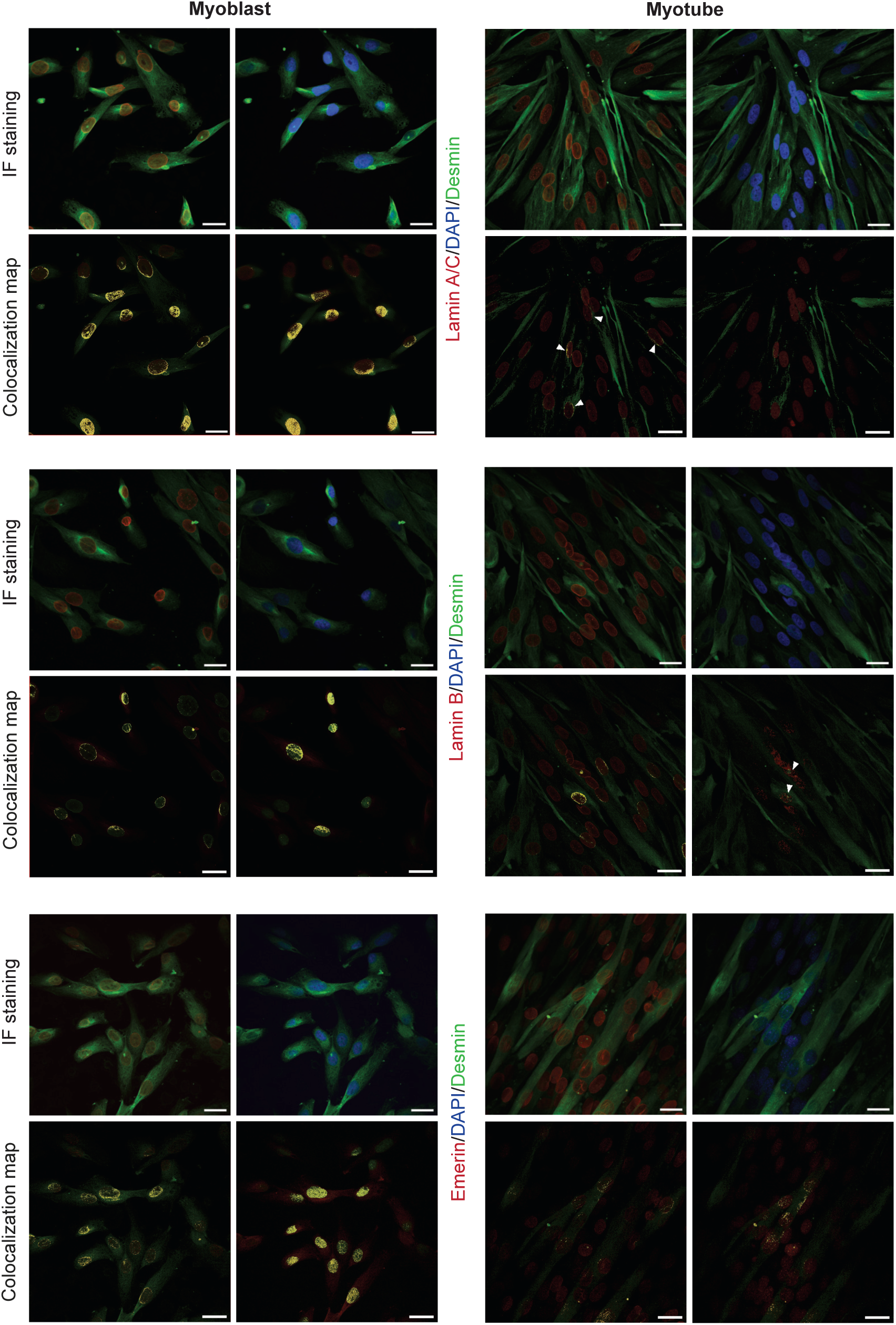
desmin is enriched at the nuclear periphery and associates with lamin A/C and emerin. Representative confocal immunofluorescence images showing desmin (green), lamin A/C, lamin B, or emerin (red), and DAPI (blue) staining in muscle cells. Corresponding fluorescence signal overlap maps generated from deconvolved images are shown . Scale bars, 25 *µ*m.

**Figure 2.**
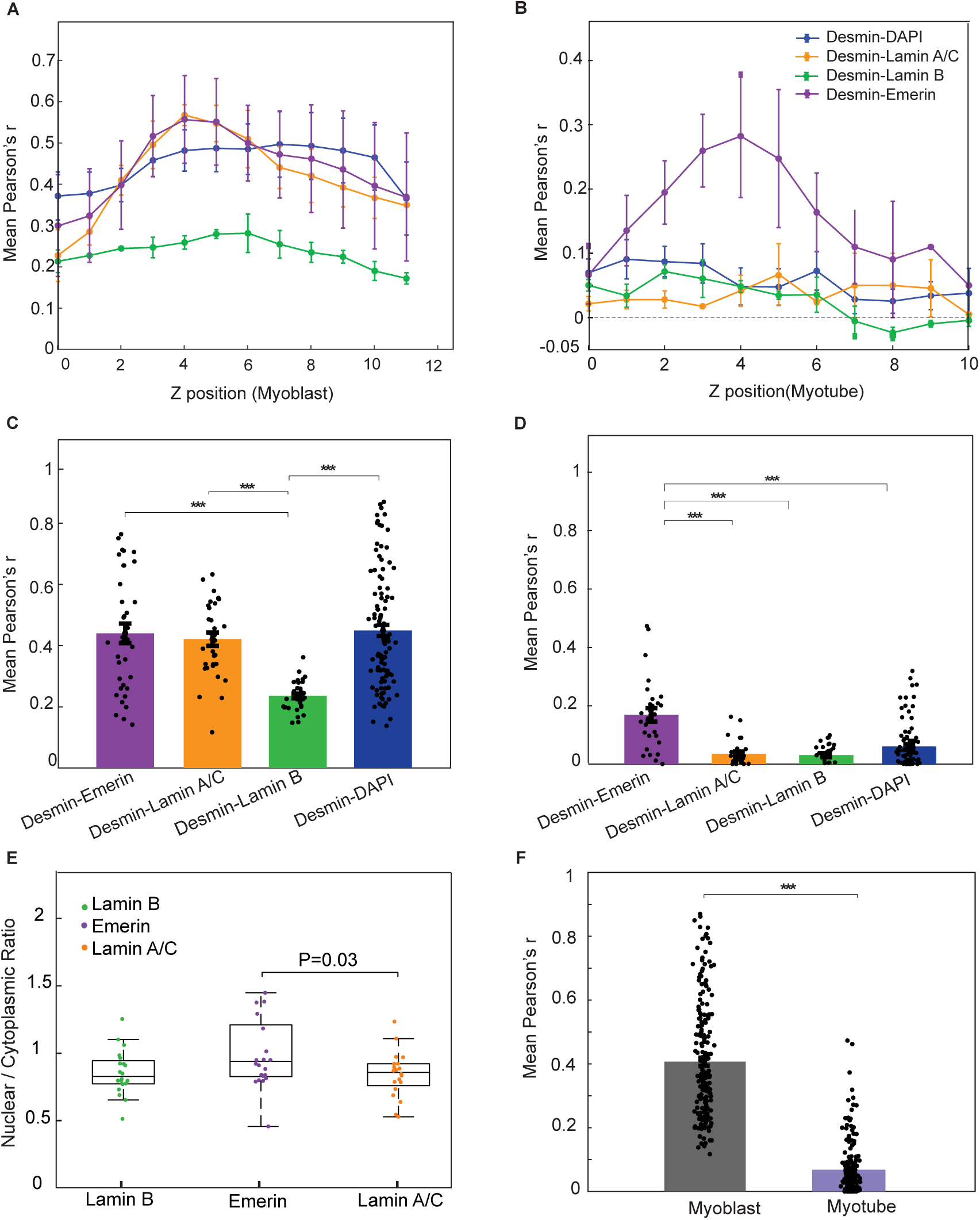
Quantitative analysis of desmin colocalization with nuclear markers during myogenic differentiation. (A) Mean Pearson’s correlation coefficients (± SEM) for desmin colocalization with DAPI, lamin A/C, Lamin B, and emerin across confocal z-stack positions in myoblasts. (B) Mean Pearson’s correlation coefficients (± SEM) for desmin colocalization with DAPI, lamin A/C, Lamin B, and emerin across confocal z-stack positions in myotubes. (C) Comparison of the overall mean Pearson’s correlation coefficients between desmin and DAPI, lamin A/C, Lamin B, and emerin in myoblasts. (D) Comparison of the overall mean Pearson’s correlation coefficients between desmin and DAPI, lamin A/C, Lamin B, and emerin in myotubes. (E) Nuclear-to-cytoplasmic (N/C) intensity ratio of desmin in Lamin B-, emerin-, and lamin A/C-positive regions in myoblasts. (F) Overall comparison of mean Pearson’s correlation coefficients between myoblasts and myotubes. Bars represent mean ± SEM, box plots indicate the median and interquartile range, and dots represent individual measurements. Statistical significance is indicated in the graphs (*\*\*\*p<* 0.001).

Quantitative analysis across z-stack sections demonstrated distinct spatial association profiles among the analyzed nuclear proteins. Pearson’s correlation coefficient analysis revealed significantly higher fluorescence signal overlap between desmin and emerin or lamin A/C compared with lamin B (Figure 1). Mean Pearson’s *r* values for desmin–emerin and desmin– lamin A/C associations peaked at intermediate z-positions (Figure 2 A–B) corresponding to the nuclear envelope, whereas desmin–lamin B association remained comparatively lower throughout the analyzed z-axis. Desmin–DAPI analyses exhibited intermediate correlation values, consistent with partial nuclear-associated signal enrichment.

### Myogenic differentiation alters the spatial association of desmin with nuclear envelope proteins

To quantitatively assess the spatial association of desmin with nuclear components during myogenic differentiation, Pearson’s correlation coefficient analyses were performed using confocal z-stack images. The distribution of colocalization across sequential optical sections demonstrated distinct association profiles between desmin and individual nuclear markers in both myoblasts and myotubes (Figure 2A,B). In myoblasts, desmin exhibited relatively high colocalization with DAPI, lamin A/C, and emerin throughout the z-stack, whereas its association with Lamin B remained consistently lower. In differentiated myotubes, overall Pearson’s correlation coefficients were reduced compared with myoblasts; however, emerin remained the nuclear marker showing the strongest association with desmin.

Comparison of the mean Pearson’s correlation coefficients confirmed these observations (Figure 2C,D). In myoblasts, desmin colocalized significantly more with DAPI, lamin A/C, and emerin than with Lamin B (*^***^p <* 0.001). Similarly, in myotubes, desmin–emerin colocalization was significantly greater than that observed for DAPI, lamin A/C, and Lamin B (*^***^p <* 0.001), indicating that the preferential association between desmin and emerin is maintained following myogenic differentiation.

To further evaluate the subcellular distribution of desmin, the nuclear-to-cytoplasmic (N/C) fluorescence intensity ratio was calculated for regions positive for lamin B, emerin, and lamin A/C (Figure 2E). Desmin associated with emerin-positive regions exhibited a significantly higher N/C ratio than lamin A/C-positive regions (P = 0.03), whereas no significant differences were detected between emerin and Lamin B or between Lamin B and lamin A/C.

Finally, comparison of the overall Pearson’s correlation coefficients between myoblasts and myotubes demonstrated a significant (*^***^p<* 0.001)reduction in desmin nuclear signals following differentiation (Figure 2F). Collectively, these findings indicate that although the overall nuclear association of desmin decreases during myogenic differentiation, its preferential spatial association with emerin is preserved, suggesting a specific interaction of desmin with components of the inner nuclear envelope.

### Targeted proteomics confirms the presence of desmin in the nuclear fraction

To biochemically validate the subcellular localization of desmin, myoblasts were fractionated into nuclear and cytoplasmic compartments. The purity of the isolated fractions was first assessed by immunoblotting using Lamin B and GAPDH as nuclear and cytoplasmic markers, respectively. Lamin B was detected exclusively in the nuclear fraction, whereas GAPDH was enriched in the cytoplasmic fraction, confirming successful fractionation (Supplementary Figure S2). Immunoblot analysis detected desmin in both the cytoplasmic and nuclear fractions, consistent with the immunofluorescence findings and indicating that a subset of desmin is associated with the nuclear compartment.

To further confirm these observations at the peptide level, targeted multiple reaction monitoring/selected reaction monitoring (MRM/SRM) mass spectrometry was performed using the desmin-specific tryptic peptide AQYETIAAK. A stable isotope-labeled AQYETIAAK peptide was first analyzed to establish the reference retention time and characteristic MRM transition profile (Figure 3A). The labeled peptide eluted at approximately 5.6 min and served as a reference for identification of the endogenous peptide.

**Figure 3.**
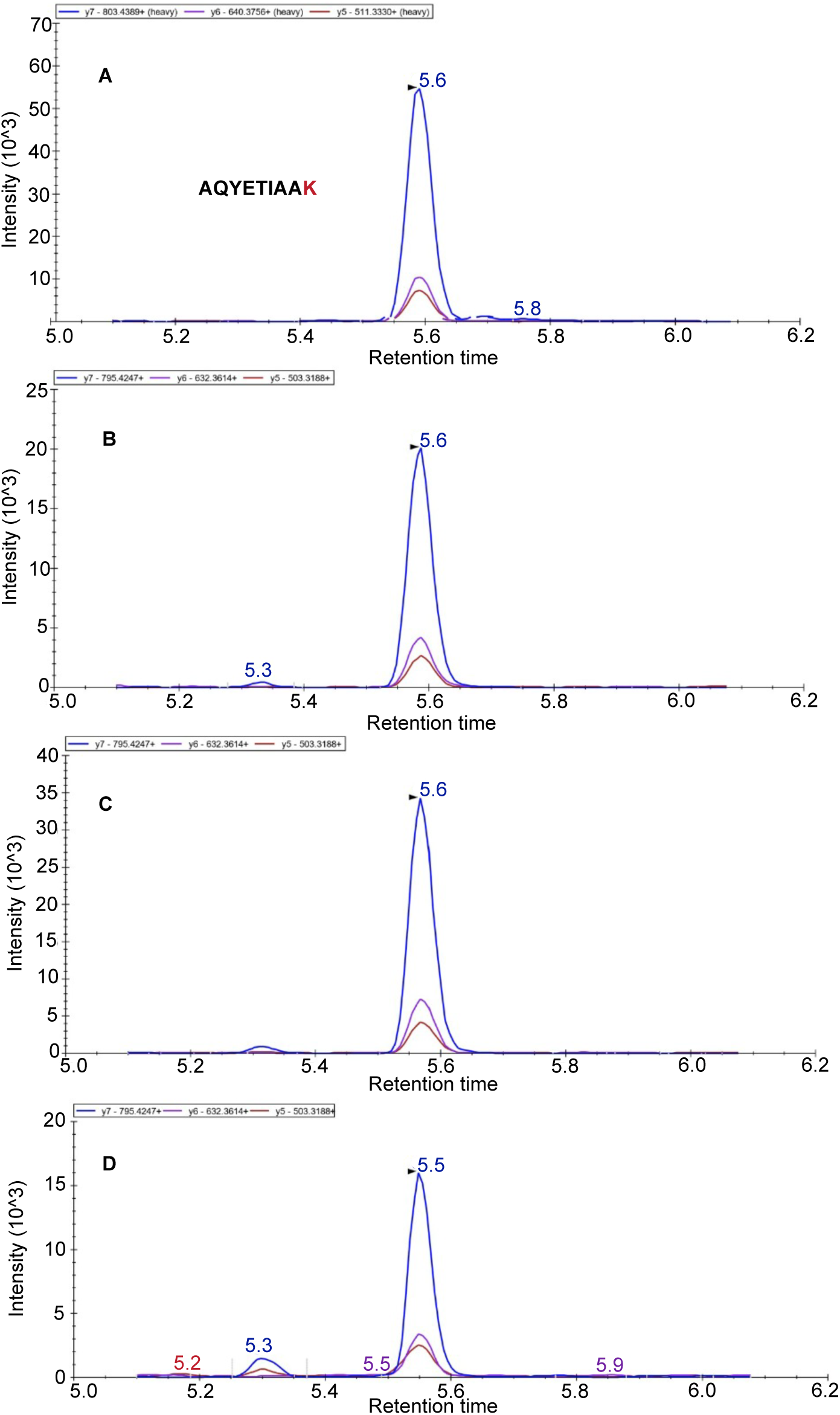
MRM-based identification of endogenous desmin using the signature peptide AQYETIAAK. (A) shows the stable isotope-labeled AQYETIAAK peptide standard, which establishes the reference retention time at approximately 5.6 min and MRM transition profile. (B), (C) and (D) show the corresponding endogenous AQYETIAAK peptide detected in myoblast day 0 total lysate, nuclear fraction, and cytosolic fraction, respectively.

To further confirm these observations, targeted proteomic analysis was performed using liquid chromatography–tandem mass spectrometry (LC–MS/MS) operated in multiple reaction monitoring (MRM) mode.

The desmin-specific tryptic peptide AQYETIAAK was selected, and its sequence specificity was verified against the UniProt database using BLAST. A stable isotope-labeled (SIL) AQYETIAAK peptide (heavy) was synthesized and used for MRM method development.

Figure 3A shows the SIL peptide detected at 5.6 min with its characteristic transition profile, which served as a reference for identification of the endogenous peptide.

The endogenous AQYETIAAK peptide was subsequently detected in total cell lysate and in the nuclear and cytoplasmic fractions isolated from day 0 myoblasts (Figure 3B–D). In all fractions, the monitored fragment ion transitions co-eluted at the expected retention time and exhibited transition patterns that matched those of the isotope-labeled peptide standard. Because AQYETIAAK is a unique desmin-derived tryptic peptide, its detection confirms the presence of desmin in both the nuclear and cytoplasmic fractions.

Together, the immunoblotting and targeted proteomic analyses independently confirm the presence of desmin within the nuclear compartment and support the imaging-based observations of nuclear desmin localization.

### Protein–protein docking predicts stable interactions between desmin and nuclear envelope proteins

To investigate whether desmin can directly associate with nuclear envelope proteins, protein–protei docking analyses were performed using desmin and the Ig-like domains of lamin B1, lamin A/C, and emerin. The top-ranked docking models predicted stable interaction interfaces for all three protein pairs (Figure 4A). In each complex, the binding interface was located along the desmin coiled-coil region, although the orientation and contact surfaces differed among the three nuclear envelope proteins.

**Figure 4.**
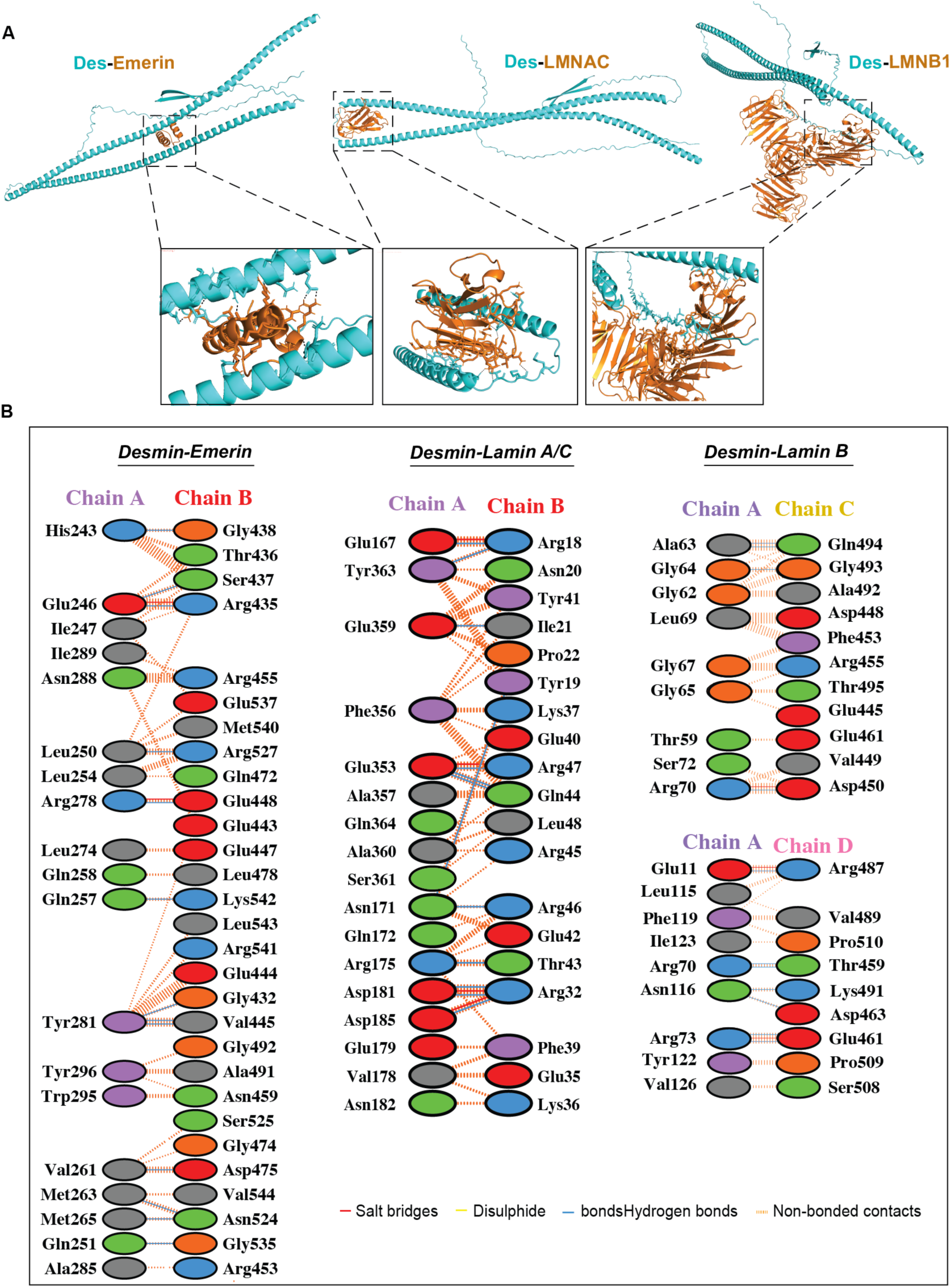
Predicted interaction interfaces between desmin and nuclear envelope proteins. (A) Three-dimensional representations of the predicted desmin–protein complexes obtained by protein–protein docking. From left to right, the desmin–emerin, desmin–lamin A/C, and desmin–lam B1 complexes are shown, respectively. In all complexes, desmin is depicted in cyan, while emerin, lamin A/C, and lamin B1 are shown in orange. Enlarged views of the predicted binding interfaces are presented below each complex. (B) Two-dimensional interaction diagrams of the corresponding protein–protein interfaces.

The predicted desmin–emerin complex positioned emerin between the two desmin *α*-helices, whereas lamin A/C and lamin B1 interacted predominantly with the terminal region of the desmin coiled-coil. Enlarged views of the docking interfaces demonstrated extensive contact surfaces involving the Ig-like domains of lamin A/C and lamin B1 as well as multiple interaction sites with emerin (Figure 4A).

Residue-level interaction analysis revealed that all three complexes were stabilized by extensive hydrogen bonds, salt bridges, and non-bonded contacts (Figure 4B). Among the predicted complexes, the desmin–lamin B1 and desmin–emerin interactions generated the most favorable docking scores and highly populated ClusPro clusters, suggesting stable interaction interfaces. The desmin–lamin A/C complex also exhibited a favorable docking configuration with an extensive interaction network. Detailed residue interactions, docking scores, cluster statistics, and Ramachandran plot analyses are provided in the Supplementary Information (Supplementary Figure S3-6).

Collectively, these computational analyses predict that desmin is capable of forming stable interactions with multiple nuclear envelope proteins, including lamin B1, lamin A/C, and emerin, supporting the experimental observations of their spatial association in myogenic cells.

## DISCUSSION

The present study provides converging evidence that endogenous desmin localizes to the nuclear compartment and preferentially associates with specific components of the nuclear envelope in human skeletal myoblasts. By integrating quantitative confocal imaging, biochemical fractionation, targeted MRM/SRM proteomics, and computational structural analyses, we demonstrate that nuclear desmin is not randomly distributed but exhibits preferential spatial association with emerin and lamin A/C. These findings extend the emerging concept that intermediate filament proteins contribute directly to nucleo-cytoskeletal communication and nuclear organization in addition to their established structural roles in the cytoplasm. Given the central role of the nuclear envelope in coordinating nuclear mechanics, chromatin organization, and mechanotransduction through interactions between lamins, LINC complex proteins, and the cytoskeleton^23–27^, our findings suggest that nuclear desmin may represent a previously underappreciated component of this regulatory network.

A major finding of this study is that endogenous desmin exhibited stronger colocalization with emerin and lamin A/C than with lamin B1, suggesting that nuclear desmin preferentially localizes to specific functional domains of the nuclear envelope. Although all three proteins are integral components of the nuclear envelope, emerin and lamin A/C are central regulators of nuclear mechanics, chromatin organization, mechanotransduction, and myogenic gene expression^28–31^. Interestingly, despite an overall reduction in nuclear desmin during myogenic differentiation, its colocalization with emerin was maintained, suggesting that this spatial association remains functionally relevant throughout myogenesis. Moreover, nuclear desmin localization was not uniform across the cell population. Only a subset of nuclei within the same imaging field exhibited detectable desmin colocalization with nuclear envelope proteins, and a similar heterogeneous distribution was observed within multinucleated myotubes, where adjacent nuclei displayed variable degrees of desmin association. This heterogeneity suggests that nuclear localization of desmin is a regulated and dynamic process rather than a constitutive feature of all myonuclei, potentially reflecting differences in nuclear functional state during myogenic differentiation. Emerin coordinates chromatin remodeling and myogenic transcription through interactions with HDAC3^32^, histone-modifying enzymes, and the LINC complex^30^, making it a plausible scaffold for a specialized nuclear desmin population during differentiation. In contrast, our previous work in zebrafish demonstrated a physical association between desmin and lamin B^33^, supporting the existence of evolutionarily conserved desmin–nuclear envelope interactions. Together, these findings suggest that nuclear desmin may engage distinct nuclear envelope partners in a context-dependent manner, with emerin and lamin A/C predominating during human myogenesis while lamin B interactions may contribute to nuclear architecture in other cellular or developmental contexts. Although direct molecular interactions remain to be established in human cells, our results further support the emerging concept that intermediate filament proteins contribute directly to nuclear organization and mechanosensitive signaling in addition to their established cytoplasmic structural functions.

Our biochemical analyses further strengthen this conclusion. Nuclear fractionation consistently detected desmin in purified nuclear fractions, and targeted MRM proteomics^34^ independently confirmed the presence of the desmin-specific peptide AQYETIAAK within the nuclear compartment. In each analyzed sample, the monitored MRM transitions were free of detectable interference, co-eluted at the expected retention time, and exhibited transition intensity patterns that closely matched those of the corresponding isotope-labeled internal standard. The concordance between endogenous and heavy-labeled peptide signals confirms the identity of AQYETIAAK and supports its reliable detection across the analyzed subcellular fractions. As AQYETIAAK is a unique desmin-derived tryptic peptide, these peptide-level data provide highly specific biochemical evidence for desmin and exclude potential ambiguity arising from the structural and sequence similarity of desmin to other type III intermediate filament proteins, such as vimentin.

Because biochemical isolation of intermediate filament proteins is technically challenging owing to their low solubility and tight association with cytoskeletal and nuclear structures, previous studies investigating nuclear intermediate filament localization have relied predominantly on microscopy-based approaches.^26,35^ The concordance between quantitative imaging, biochemical fractionation, and peptide-level proteomic identification therefore provides robust and complementary evidence that endogenous desmin is present within the nuclear compartment, substantially reducing the likelihood that our observations result from imaging artifacts or fractionation contamination. These findings are consistent with our recent demonstration that desmin undergoes active nuclear import through an importin *α/β*-dependent mechanism mediated by two functional nuclear localization signals^20^, and extend this work by providing biochemical evidence for endogenous nuclear desmin together with its preferential association with specific components of the nuclear envelope.

To gain structural insight into the observed colocalization patterns, we performed protein–protei docking analyses between desmin and the nuclear envelope proteins lamin B1, lamin A/C, and emerin. All three proteins generated structurally plausible binding interfaces with desmin through complementary hydrogen-bonding, electrostatic, and hydrophobic interactions, supporting the possibility that desmin is capable of engaging multiple components of the nuclear envelope. Notably, both lamin B1 and lamin A/C were predicted to interact predominantly through their C-terminal Ig-like domains, which are well-established protein–protein interaction modules involved in the recruitment and stabilization of nuclear envelope proteins.^36,37^ In contrast, emerin adopted a distinct binding mode, occupying the groove between the two coiled-coil *α*-helices of desmin. This observation is consistent with the intrinsically disordered nature of emerin, which enables it to interact with numerous structural and regulatory partners, including lamins, actin, barrier-to-autointegration factor (BAF), and LINC complex components, thereby functioning as a dynamic scaffold at the inner nuclear membrane.^1,30^ The distinct interaction interfaces predicted for emerin, lamin A/C, and lamin B1 are also consistent with the structural diversity of these nuclear envelope proteins. While the lamin proteins contain compact Ig-like domains that provide well-defined protein interaction surfaces, emerin possesses an extended intrinsically disordered architecture in which only the LEM domain adopts a stable globular fold. Consequently, desmin is unlikely to recognize a single conserved binding motif and instead appears capable of adapting to structurally different molecular surfaces presented by individual nuclear envelope proteins. Comparison of the docking solutions further supported these structural differences. The desmin–emerin complex produced the largest docking cluster (165 members) together with one of the most favorable docking scores (center energy: *—*789.8), whereas the desmin–lamin B1 complex also exhibited highly favorable docking parameters (center energy: *—*786.2; cluster size: 72 members). The desmin–lamin A/C complex generated a slightly less favorable docking solution (center energy: *—*707.5; cluster size: 68 members) but retained an extensive interaction interface and acceptable stereochemical quality. Residue-level interaction analyses further indicated that the predicted interfaces were stabilized predominantly by complementary hydrogen-bonding and electrostatic interaction networks, while hydrophobic contacts contributed to interface packing. Despite differences in the interacting residues among emerin, lamin A/C, and lamin B1, all three complexes exhibited chemically complementary interaction surfaces, supporting the structural plausibility of the predicted binding modes. Although docking analyses alone cannot establish direct molecular interactions, the predicted interfaces were consistent with our confocal colocalization and biochemical fractionation data, providing a plausible structural framework through which desmin may associate with multiple nuclear envelope proteins. Together, these findings support a model in which nuclear desmin contributes to the organization of a mechanically responsive nuclear envelope rather than interacting with a single nuclear binding partner.

The present findings also have important implications for understanding desmin-related muscle disease. We previously identified a pathogenic *DES* splice-site mutation associated with limb-girdle muscular dystrophy R2^21^that impairs myogenic differentiation and markedly reduces myotube formation.^22^ The present study extends these observations by suggesting that disruption of desmin interactions with the nuclear envelope may represent an additional mechanism contributing to disease pathogenesis. Our combined imaging, biochemical, proteomic, and structural analyses indicate that nuclear desmin preferentially associates with emerin and lamin A/C while retaining the capacity to interact with lamin B1, supporting a role in maintaining nuclear architecture and mechanosensitive signaling. Perturbation of these interactions could therefore compromise chromatin organization, mechanotransduction, and transcriptional programs required for efficient muscle differentiation, consistent with the established functions of emerin and A-type lamins in myogenesis and genome organization. Collectively, our findings support the concept that desmin-related muscle disease may arise not only from disruption of the cytoplasmic intermediate filament network but also from impaired nucleo-cytoskeletal communication and nuclear envelope function.

Rather than functioning as an isolated nuclear protein, our findings suggest that desmin forms part of a dynamic nuclear envelope interaction network that links the intermediate filament cytoskeleton to nuclear architecture. The preferential association of desmin with emerin and lamin A/C, together with its conserved capacity to associate with lamin B, supports the concept that nuclear desmin engages distinct nuclear envelope partners in a context-dependent manner to coordinate structural and regulatory functions during myogenesis. This work therefore expands the current view of desmin biology beyond its established cytoplasmic role and identifies the nuclear envelope as a previously unrecognized functional platform for desmin. Elucidating how these interactions influence chromatin organization, mechanosensitive signaling, and transcriptional regulation may provide new insights into the molecular basis of desmin-related myopathies and other disorders associated with impaired nucleo-cytoskeletal communication.

### Limitations of the study

Several limitations should be considered when interpreting the present findings. First, variability in protein concentrations and cell yields following nuclear fractionation posed challenges for data normalization, particularly because endogenous nuclear desmin represents a low-abundance protein population. In addition, the absence of technical replicates in the targeted proteomic analyses limited the statistical robustness of peptide quantification. The limited recovery of nuclear protein also precluded complementary biochemical interaction studies, such as coimmunoprecipitation or affinity-based pull-down assays, which would have provided direct experimental validation of the predicted desmin–nuclear envelope protein interactions in human skeletal muscle cells. The present study focused on three representative nuclear envelope proteins—lamin B1, lamin A/C, and emerin. Future studies should determine whether nuclear desmin participates in a broader interaction network involving additional structural and regulatory proteins within the nuclear envelope. Finally, the present study was performed in cultured human skeletal muscle cells; therefore, the functional significance of nuclear desmin interactions during muscle development and in desmin-related myopathies remains to be established in vivo.

## RESOURCE AVAILABILITY

### Lead contact

Requests for further information and resources should be directed to and will be fulfilled by the lead contact, Nilufer Duz.

### Materials availability

This study did not generate new materials.

### Data and code availability

- Any additional information required to reanalyze the data reported in this paper is available from the lead contact upon request.

## ACKNOWLEDGMENTS

This work was supported by TÜBİTAK through grants 123Z114 and 224S559, and by EMBO through grant 9337. This project was also partially funded by METU grants GAP-103-2023-11139 and ADEP-103-2023-11251. The authors gratefully acknowledge all members of the Q-OmicS Lab, with special thanks to Ceren Kul for her support during the analysis. We also thank Shimadzu Corporation for providing the LCMS-8060 instrument, which significantly contributed to the success of this study. The authors additionally thank Dr. Carlijn Bouten and Dr. Rob Driessen for their valuable support.

## AUTHOR CONTRIBUTIONS

Conceptualization, N.D. and P.D. ; methodology, N.D., A.A., and S.S., M.K.; investigation, N.D. and S.O.; writing-–original draft, N.D. and S.O and A.A.; writing-–review & editing, N.D. and P.D.; funding acquisition, N.D, S.O. and P.D.; resources, N.D and P.D.; supervision, N.D., S.O. and P.D.

## DECLARATION OF GENERATIVE AI AND AI-ASSISTED TECHNOLOGIES

During the preparation of this work, the author(s) used [ChatGPT (version 4)] in order to [typo and grammar check.]. After using this tool or service, the author(s) reviewed and edited the content as needed and take(s) full responsibility for the content of the publication.

## STAR METHODS

### Resource availability

#### Lead contact

Further information and requests for resources and reagents should be directed to and will be fulfilled by the lead contact.

#### Materials availability

This study did not generate new unique reagents.

#### Data and code availability

The datasets and MATLAB codes generated and analyzed during this study are available from the corresponding author upon reasonable request.

### Experimental model and subject details

#### Cell culture models

Immortalized human skeletal muscle myoblasts (ABM, Cat# T0033, RRID: CVCL VG47) were used as the experimental model in this study. Cells were maintained under standard culture conditions and differentiated into multinucleated myotubes for differentiation experiments as described below.

### Method details

#### Cell culture

Immortalized human skeletal muscle myoblasts (ABM, Cat# T0033, RRID: CVCL VG47) were cultured in high-glucose (4.5 g/L) Dulbecco’s Modified Eagle Medium (DMEM) supplemented with stable L-glutamine (Gibco, Thermo Fisher Scientific, Waltham, MA, USA), 20% fetal bovine serum (FBS), 50 *µ*g/mL fetuin, 10 ng/mL epidermal growth factor (EGF), 1 ng/mL basic fibroblast growth factor (bFGF), 10 *µ*g/mL insulin, 0.4 *µ*g/mL dexamethasone, and 1% gentamicin/amphotericin B. Cells were maintained at 37*^○^*C in a humidified incubator with 5% CO_2_.

Myogenic differentiation was induced when cultures reached greater than 90% confluence by replacing the growth medium with differentiation medium consisting of high-glucose (4.5 g/L) DMEM supplemented with stable L-glutamine, 2% horse serum, and 10 *µ*g/mL recombinant human insulin. Cells were maintained under differentiation conditions for 6 days at 37*^○^*C in a humidified atmosphere containing 5% CO_2_ before subsequent analyses.

#### Immunofluorescence staining

Cells were fixed with 4% paraformaldehyde for 10 minutes and subsequently permeabilized and blocked using 2% FBS in PBS containing 0.1% Triton X-100 for 1 hour at room temperature. Samples were then incubated overnight at 4*^○^*C with primary antibodies diluted in PBS. The following primary antibodies were used: Desmin (Sigma-Aldrich, D8281, 1:100), Lamin B1 (MA1-06103, Invitrogen, 1:100),lamin A/C (clone 131C3, ab8984, Abcam, 1:100), and Emerin (clone 4G5, NCL-EMERIN, Leica Biosystems, 1:100). Following three washes with 1*×* PBS, cells were incubated with secondary antibodies, goat anti-mouse IgG (H+L) Alexa Fluor 488 (A11001, Invitrogen, 1:1000) and goat anti-rabbit IgG Alexa Fluor 555 (A21428, Invitrogen, 1:1000), diluted in PBS. After three additional washes with 1*×* PBS, nuclei were stained with DAPI (D9542, Sigma-Aldrich, 1:500) for 5 minutes. Samples were mounted using Mowiol (Sigma-Aldrich).

Images were acquired using a Zeiss LSM 880 laser scanning confocal microscope equipped with a 63*×*/1.4 Apochromat Oil DIC objective (Carl Zeiss, Germany). Colocalization analyses were performed using Pixel-based analyses (quantification of overlap between fluorescence intensity distributions in different channels). Following deconvolution, colocalization analyses were performed using Huygens Professional software version 23.04 (Scientific Volume Imaging, The Netherlands; http://svi.nl). Samples were prepared in three independent experiments, and images from three distinct fields were acquired for each sample. For nuclear-to-cytoplasmic (N/C) fluorescence intensity analysis, the same cells used for the colocalization analysis were evaluated. Nuclear regions of interest (ROIs) were manually delineated based on DAPI staining, whereas the entire desmin-positive cytoplasm of each corresponding cell, excluding the nucleus, was manually outlined in Fiji software (RRID: SCR 002285). Mean desmin fluorescence intensity was measured separately within the nuclear and cytoplasmic ROIs, and the N/C ratio was calculated by dividing the mean nuclear fluorescence intensity by the mean cytoplasmic fluorescence intensity. All images were acquired using identical microscope settings, and ROI selection and fluorescence intensity measurements were performed using identical analysis parameters for all samples.

#### Subcellular fractionation and immunoblotting

Nuclear and cytosolic fractions were isolated from muscle progenitor cells using an optimized fractionation protocol adapted from Dimauro et al. (2012). Briefly, scraped cells were washed with cold PBS and collected by centrifugation at 200 *×* g for 7 minutes. Cell pellets were resuspended in 300 *µ*l STM buffer containing 250 mM sucrose, 50 mM Tris-HCl (pH 7.4), 5 mM MgCl_2_, and protease/phosphatase inhibitor cocktails (Sigma-Aldrich, Poole, UK). Homogenization was performed on ice for 1 minute using a tightly fitting Teflon pestle attached to a Potter S homogenizer (İLDAM, Ankara, Turkey). Homogenates were incubated on ice for 30 minutes, vortexed for 15 seconds at maximum speed, and centrifuged at 800 *×* g for 15 minutes. The resulting pellet was collected as the nuclear fraction, whereas the supernatant was designated as the cytoplasmic fraction. Further steps included resuspending the pellet in 300 *µ*l STM buffer, vortexing for 15 seconds, and centrifugation at 500 *×* g for 15 minutes. The resulting pellet was resuspended in 200 *µ*l NET buffer containing 20 mM HEPES (pH 7.9), 1.5 mM MgCl_2_, 0.5 M NaCl, 0.2 mM EDTA, 20% glycerol, 1% Triton X-100, and protease/phosphatase inhibitor cocktails. The mixture was vortexed for 15 seconds at maximum speed and incubated on ice for 30 minutes. Nuclei were subsequently lysed by sonication using three cycles at 50% amplitude for 20 seconds each while maintaining samples on ice. Sonication was performed using a Vibra-Cell probe sonicator (model VCX 130) equipped with a CV 18 probe (Sonics & Materials, Newton, CT, USA). The lysate was centrifuged at 9,000 *×* g for 30 minutes at 4*^○^*C, and the resulting supernatant was collected as the nuclear fraction.

The cytosolic fraction was further centrifuged at 11,000 *×* g for 10 minutes. The resulting supernatant, containing cytosolic and microsomal fractions, was precipitated with cold 100% acetone at *—*20*^○^*C for at least 1 hour. Following centrifugation at 12,000 *×* g for 5 minutes, the pellet was resuspended in 100–300 *µ*l STM buffer and designated as the cytosolic fraction.

To extract total proteins, cells were rinsed with cold PBS and lysed in RIPA buffer (Sigma-Aldrich, St. Louis, MO, USA) supplemented with 1*×* protease inhibitor cocktail (Roche, Basel, Switzerland) and 1 mM sodium orthovanadate. Samples were sonicated six times at 50% amplitude for 20 seconds each while maintained on ice. Cell debris was removed by centrifugation, and protein concentrations were determined using the BCA Protein Assay (Pierce) according to the manufacturer’s instructions. Absorbance at 562 nm was measured using a SpectraMax M2 Microplate Reader (Molecular Devices, San Jose, CA, USA).

Equal amounts of protein (25 *µ*g) were loaded onto SDS-PAGE gels. Western blot analysis was performed using the following primary antibodies: Lamin B (nuclear marker; ab901, Abcam, 1:1000), glyceraldehyde 3-phosphate dehydrogenase (GAPDH; cytosolic marker; G9295, Sigma-Aldrich, 1:1000), and Desmin (D8281, Sigma-Aldrich, 1:200). Membranes were subsequently incubated with horseradish peroxidase (HRP)-conjugated anti-mouse secondary antibody (12-349, Sigma-Aldrich, 1:3000), anti-chicken secondary antibody (A16054, Thermo Fisher Scientific, 1:5000), and HRP-conjugated anti-rabbit secondary antibody (ab6721, Abcam, 1:1000). Chemiluminescence detection was performed using the GeneGnome 5 imaging system (Syngene, Bangalore, India). Samples were prepared in three independent experiments.

#### Protein precipitation and tryptic digestion

Proteins were precipitated using four volumes of 80% (v/v) cold acetone and incubated for 18 h at *—*20*^○^*C. Following precipitation, samples were centrifuged at 12,000 *×* g for 20 min, and the supernatant was discarded. Protein pellets were dried for 1 h at 45*^○^*C using a vacuum evaporator (Concentrator 5301, Eppendorf) and subsequently resuspended in 100 *µ*L of 50 mM ammonium bicarbonate (ABC; Sigma-Aldrich).

For reduction, 10 mM dithiothreitol (DTT; Sigma-Aldrich) prepared in 50 mM ABC was added, and samples were incubated for 5 min at 95*^○^*C. Alkylation was performed using 15 mM iodoacetamide (IAA; Sigma-Aldrich) in 50 mM ABC for 20 min at room temperature in the dark. Proteolytic digestion was carried out overnight at 37*^○^*C using Trypsin/LysC protease (Promega) at a 1:50 enzyme-to-protein ratio.^39^

Digestion was quenched with 0.1% LC-MS-grade formic acid (Merck), and samples were dried in a vacuum evaporator to remove residual solvent. Peptides were reconstituted in 100 *µ*L of 0.1% LC-MS-grade formic acid prepared in LC-MS-grade water (Merck) prior to LC-MRM/MS analysis.

Protein concentrations of the total protein, nuclear, and cytoplasmic fractions were determined using the bicinchoninic acid (BCA) protein assay (Thermo Fisher Scientific) before proteolytic digestion. The endogenous AQYETIAAK peptide abundance was normalized to the total protein content and expressed as nmol desmin per *µ*g of total protein. Protein concentrations and the calculated amount of endogenous AQYETIAAK peptide in 1 *µ*g of total protein are summarized in Supplementary Table S1.

#### Liquid chromatography and triple quadrupole mass spectrometry

A unique Desmin-derived AQYETIAAK peptide (UniProt ID: P17661) was selected for targeted protein quantification. The stable isotope-labeled (SIL) form of AQYETIAAK containing a C-terminal heavy lysine residue (^13^C_6_ and ^15^N_2_) was synthesized by JPT Peptide Technologies (Berlin, Germany). Lyophilized SIL peptide was resuspended in a solution containing 80% 0.1 M ammonium bicarbonate (ABC) and 20% LC-MS-grade acetonitrile (ACN; Merck) according to the manufacturer’s instructions. Peptide purity was assessed by high-performance liquid chromatography (HPLC; Agilent) and determined to be 44% by PeptiTeam (Ankara, Türkiye). Mass validation was performed using LC-QMS (Agilent).

Tryptic peptides were separated using an AdvanceBio Peptide Mapping column (2.1 *×* 150 mm, 2.7 *µ*m; Agilent) maintained at 40*^○^*C. Chromatographic separation was performed using solvent A (LC-MS-grade water containing 0.1% formic acid) and solvent B (acetonitrile containing 0.1% formic acid) at a flow rate of 0.3 mL/min. The gradient elution profile was as follows: 95% A/5% B for 4 min, 85% A/15% B for 16 min, 75% A/25% B for 10 min, 45% A/55% B for 5 min, 20% A/80% B for 5 min, followed by re-equilibration at 95% A/5% B for 10 min.

Mass spectrometric analysis was performed using a Shimadzu LCMS-8060 triple quadrupole mass spectrometer equipped with an electrospray ionization (ESI) source. Instrument parameters were set as follows: nebulizing gas flow, 3 L/min; heating gas flow, 10 L/min; interface temperature, 300*^○^*C; desolvation temperature, 526*^○^*C; desolvation line temperature, 250*^○^*C; heat block temperature, 400*^○^*C; drying gas flow, 10 L/min; and oven temperature, 40*^○^*C.

Method optimization was performed using SIL peptide standards. Peptide identification and quantification were carried out using three optimized transitions corresponding to y7, y6, and y5 fragment ions with mass-to-charge (m/z) ratios of 803.44, 640.38, and 511.33, respectively. Matrix-matched calibration curves were generated using seven concentration points ranging from 0.2 to 10 nmol/*µ*L. Endogenous peptide analysis was performed with SIL peptide spike-in at a concentration of 2.09 *µ*mol/*µ*L per sample, and the injection volume was set to 10 *µ*L.

#### LC-MRM/MS data analysis

Data processing, peak picking, and peak integration were performed using Skyline software (version 24.1; Washington University, USA). Transitions were filtered to minimize matrix effects and signal interference. Quantification was performed using the highest interference-free peak area for each transition. The ratio of endogenous (light) peptide peak area to stable isotope-labeled (heavy) peptide peak area was calculated for each sample. Endogenous peptide abundance was determined using the following equation:^40^

Amount of endogenous peptide (nmol) = Light-to-heavy ratio *×* SIL peptide amount (nmol) Calculated endogenous peptide amounts were normalized to 1 *µ*g of total protein extract.

#### Protein–protein docking analysis

The protein structures used in the docking studies were obtained from the RCSB Protein Data Bank (PDB) and AlphaFold Protein Structure Database. The crystal structure of the human lamin B1 immunoglobulin-like (Ig-like) domain (PDB ID: 7DTG)^41^ and the crystal structure of the human lamin A/C Ig-like domain (PDB ID: 1IFR)^36^, both of which are known to participate in protein–protein interactions within the nuclear lamina, were retrieved from the RCSB Protein Data Bank. Due to the limited availability of experimentally determined structural data for desmin, the AlphaFold-predicted human desmin structure (AlphaFold ID: AF-AFP17661-F1)^42^ was employed as a model. Prior to docking studies, the structure was refined using Swiss-Model. The resulting model exhibited a GMQE (Global Model Quality Estimation) score of 0.77, indicating acceptable structural reliability. Since no complete high-resolution experimental structure of human emerin is currently available, the AlphaFold-predicted human emerin structure (UniProt ID: Q14126) was used for protein–protein docking studies. Because substantial regions of emerin are intrinsically disordered, only the relatively structured N-terminal segment comprising residues 8–48 was retained for docking analyses. This region was selected in order to reduce conformational ambiguity and improve the structural reliability of the docking calculations. Protein structures were prepared using the Protein Preparation Wizard module of Schrödinger Suite version 12.0.6 (Release 2024-2). Protein–protein docking calculations were performed using the ClusPro^43^. Three independent docking systems were evaluated: desmin–lamin B1, desmin–lamin A/C, and desmin–emerin. The resulting protein complexes were subsequently analyzed using PDBsum and PyMOL (The PyMOL Molecular Graphics System, Version 4.5, Schrödinger, LLC) software packages to characterize intermolecular contacts, interaction interfaces, and structural features of the predicted complexes. The stereochemical quality of the docked complexes was further evaluated through Ramachandran plot analysis.

#### Quantification and statistical analysis

Statistical analyses and graphical visualizations were performed using MATLAB (MathWorks, Natick, MA, USA). Pearson’s correlation coefficients (Pearson’s r) were calculated for pixel-based colocalization analyses between Desmin and Lamin A/C, Lamin B, emerin, or DAPI signals following image deconvolution and colocalization analysis in Huygens Professional software version 23.04 (Scientific Volume Imaging, The Netherlands). Mean Pearson’s correlation coefficients were calculated from all optical sections acquired at 1 *µ*m intervals from three randomly selected fields in each of three independent biological experiments. Depending on cell density, each field contained a variable number of cells, and all cells within each field were included in the analysis.

Group comparisons were performed using one-way analysis of variance (ANOVA) followed by post hoc multiple comparison analysis. Statistical significance was defined as *p<* 0.05. Data are presented as mean *±* SEM.

